# The 15-min (Sub)Cellular Proteome

**DOI:** 10.1101/2024.02.15.580399

**Authors:** Bowen Shen, Leena R. Pade, Peter Nemes

## Abstract

Single-cell mass spectrometry (MS) opens a proteomic window onto the inner workings of cells. Here, we report the discovery characterization of the subcellular proteome of single, identified embryonic cells in record speed and molecular coverage. We integrated subcellular capillary microsampling, fast capillary electrophoresis (CE), high-efficiency nano-flow electrospray ionization, and orbitrap tandem MS. In proof-of-principle tests, we found shorter separation times to hinder proteome detection using DDA, but not DIA. Within a 15-min effective separation window, CE data-independent acquisition (DIA) was able to identify 1,161 proteins from single HeLa-cell-equivalent (∼200 pg) proteome digests vs. 401 proteins by the reference data-dependent acquisition (DDA) on the same platform. The approach measured 1,242 proteins from subcellular niches in an identified cell in the live *Xenopus laevis* (frog) embryo, including many canonical components of organelles. CE-MS with DIA enables fast, sensitive, and deep profiling of the (sub)cellular proteome, expanding the bioanalytical toolbox of cell biology.

**Authorship Contributions:** P.N. and B.S. designed the study. L.R.P. collected the *X. laevis* cell aspirates. B.S. prepared and measured the samples. B.S. and P.N. analyzed the data and interpreted the results. P.N. and B.S. wrote the manuscript. All the authors commented on the manuscript.

## INTRODUCTION

Inter- and intra-cellular molecular differences orchestrate important biological states and functions, but the identity and roles of the protein building blocks have yet to be fully understood. A key limitation has been due to a lack of technologies capable of sufficient sensitivity to characterize the proteome of the cell or its components from the trace amounts of materials contained within. Super-resolution light^[1]^ and cryo-electron microscopies^[2]^ have expanded our view of cellular processes, albeit with constraints to a few molecular targets at a time. While a lack of amplification to the whole proteome fundamentally thwarts comprehensive molecular coverage, mass spectrometry (MS) proteomics has recently gained sensitivity to individual cells (reviewed in Refs. ^[3]^), with community-wide standards advocating for rigor and reproducibility^[4]^. This nascent field of single-cell MS has already begun categorizing previously unknown proteomic differences between cells, testing hypotheses, and making biological discoveries, for example, in embryonic cells in live embryos^[5]^ and culture^[6]^, hepatocytes^[7]^, and neurons in the mouse^[8]^. Single-cell MS technologies promise transformative potentials in cell biology and health research, provided current challenges can be addressed in proteome coverage, analysis speed, and scalability.

Single-cell MS bridges technical innovations from many laboratories. Nearly 20 years after globulin analysis pioneered the prospect of targeted single-cell proteomics using capillary electrophoresis (CE) and Fourier transform ion cyclotron resonance,^[9]^ microdissection and CE electrospray ionization (ESI) MS paved the beginning of discovery^[10]^ and multiplexing quantitative proteomics^[11]^ on single cells using benchtop orbitrap^[11]^ and time-of-flight^[10]^ mass spectrometers. Concerted efforts soon followed to streamline detection sensitivity through refining sample collection to capillaries^[12]^ and cell sorting^[13]^ as well as processing of limited proteomes using specialized micro-vials^[11, 12b]^, microwell plates (e.g., SOP^[13b]^ and mPOP^[13c]^), and recently nanofabricated surfaces (e.g., nanoPOTS^[13a]^). As a consequence, CE-MS uncovered 1,709 proteins among single identified cells in *X. laevis* embryos^[11]^ and detected ∼744 proteins from a single HeLa-cell-equivalent proteome amounts using orbitrap HRMS.^[14]^ NanoPOTS sample processing improved sensitivity to more than 1,000 proteins from single HeLa cells on an ultra-low-flow nano chromatograph (nanoLC).^[15]^ SCoPE leveraged specialized barcoding and nanoLC-HRMS to 3,042 proteins among 1,490 single macrophages.^[16]^ Ion filtering based on high-field asymmetric waveform ion mobility spectrometry (FAIMS) reduced spectral complexity and ion load, furthering identifications to ∼1,100 proteins per single HeLa cells.^[17]^ For proteome amounts in mammalian cells, these workflows routine detect 1,000–2,000 proteins by nanoLC-MS^[15, 18]^ and 350–750 proteins by CE-MS^[14, 19]^.

The next evolution in single-cell proteomics calls for improved detection sensitivity, speed, and scalability. With automation and isobaric barcoding, nanoLC enabled concurrent exploration of 3,042 proteins from 1,490 cells in 10 days of total nanoLC instrument time^[16b]^. Similarly, fast chromatography within 7.2-min (5 min gradient) was able to analyze 200 samples in a day, identifying ∼1,000 proteins from 5 ng of HeLa proteome digests on a FAIMS orbitrap mass spectrometer.^[20]^ To complement these technical milestones via nanoLC, we here report a second-generation CE-MS-based technology that combines fast analysis, deep proteomics, and scalability to the subcellular regime. With a capability to achieve trace-level, 30-zmol (18,000 copies) sensitivity^[14, 21]^, exquisite separation resolution^[11]^, and scalability across broad spatial and temporal dimensions^[22]^, CE-MS has been emerging for single-cell proteomics (reviewed in Ref. ^[23]^). Most recently, CE-MS enabled *in vivo* single-cell dual proteo-metabolomics on identified cells in live *X. laevis* embryos^[12a]^ and measured 157 proteins in single neuronal somas in the mouse brain through patch-clamp proteomics ^[8]^. Development of a CE-MS with faster and deeper proteome coverage complements current efforts in nanoLC-MS to strengthen our basic understanding of the cell during states of health and disease.

## RESULTS AND DISCUSSION

### We sought to improve CE-MS sensitivity

Our experimental approach is presented in **Figure 1**. We chose the commercial HeLa proteome digest to develop and test the technology as it could be sourced reproducibly from reputable vendors. Afterward, we extended the workflow to enable subcellular proteomics in the animal-dorsal midline (called D11) cell in the 16-cell *Xenopus laevis* embryo, which are fated to reproducibly form neural tissues.^[24]^ Building on our previous work, capillary microsampling was scaled to aspirate ∼10 nL, or ∼11% of the total volume of the cell *in vivo*. Previously, one-to-two hours of separations identified ∼800 proteins from 5 ng of *Xenopus* proteome by CE-MS^[12]^ and ∼1,650 proteins from 40 ng of *Xenopus* proteome by LC-MS^[5c]^. These studies provided a reference to benchmark performance for our current study. We reasoned that contemporary single-cell proteomics would benefit from deeper molecular coverage and higher analytical throughput to allow for higher statistical power and investigations on larger cell populations. Here, we set our milestones at doubling the proteome coverage and achieving ∼4–8-times higher high-throughput by enabling CE-MS proteomics in less than 15 min of instrumental analysis.

**Figure 1.**
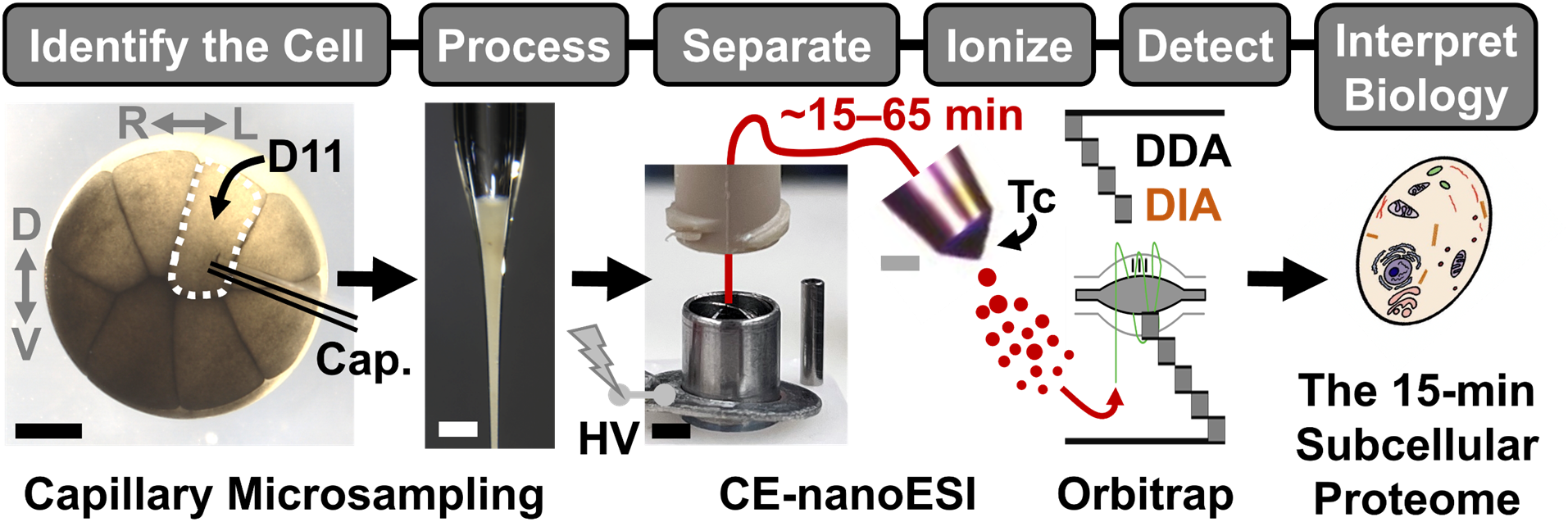
Our experimental approach to enabling 15-min subcellular proteomics using capillary electrophoresis (CE) mass spectrometry (MS). The approach was developed and validated using the commercial HeLa proteome digest. For demonstration, a subcellular niche was aspirated using a fabricated microcapillary (Cap.) from the dorsal-animal midline (called D11) cell in the 16-cell live *X. laevis* embryo. The collected proteome was processed using standard vials for bottom-up proteomics. A ∼5 ng portion of the proteome digest was electrophoresed over ∼15- to 65-min by varying the CE potential (high voltage, HV) and ionized in an electrokinetically pumped CE nano-flow ESI interface operated in the cone-jet regime (Taylor cone, Tc shown). Peptide ions were detected on a quadrupole orbitrap tandem mass spectrometer executing data-dependent acquisition (DDA) or data-independent acquisition (DIA). The resulting data informed gene ontology (GO) annotation of cellular organization and molecular roles. Scale bars, 250 μm (black), 300 μm (white).

Our first objective was to deepen the detectable proteome. As in most MS-based bottom-up proteomics, the coverage of the single-cell proteome in this work (**Fig. 1**) was determined by the number of different proteotypic peptides that could be sequenced to high quality during cycles of survey scans (MS^1^) and tandem transitions (MS/MS). We recently reported narrow, few-seconds-wide electrophoretic peaks to challenge proteome coverage by exhausting the MS– MS/MS duty cycle, which was partially remedied by a designer data acquisition method employing a DDA ladder^[19]^. However, optimization on the experimental parameter space, including the detected mass-to-charge (m/z) range to the HeLa proteome digest, did not yield additional benefits (**Fig. 2A**). A maximum of ∼1,160 ± 31 proteins (1,541 cumulative) were identifiable from 10 ng of HeLa proteome digest (*m/z* 350–900) in technical triplicates. The detected proteins are tabulated in the electronic Supplementary Information (**SI**) **Table 1** (**Table S1**). Although these results compare favorably to ∼850–1,600 proteins that were identified from comparable proteome amounts on nanoLC with comparable mass spectrometers^[25]^, a saturation of the duty cycle indicated a fundamental limitation to furthering coverage through optimization via CE-MS.

**Figure 2.**
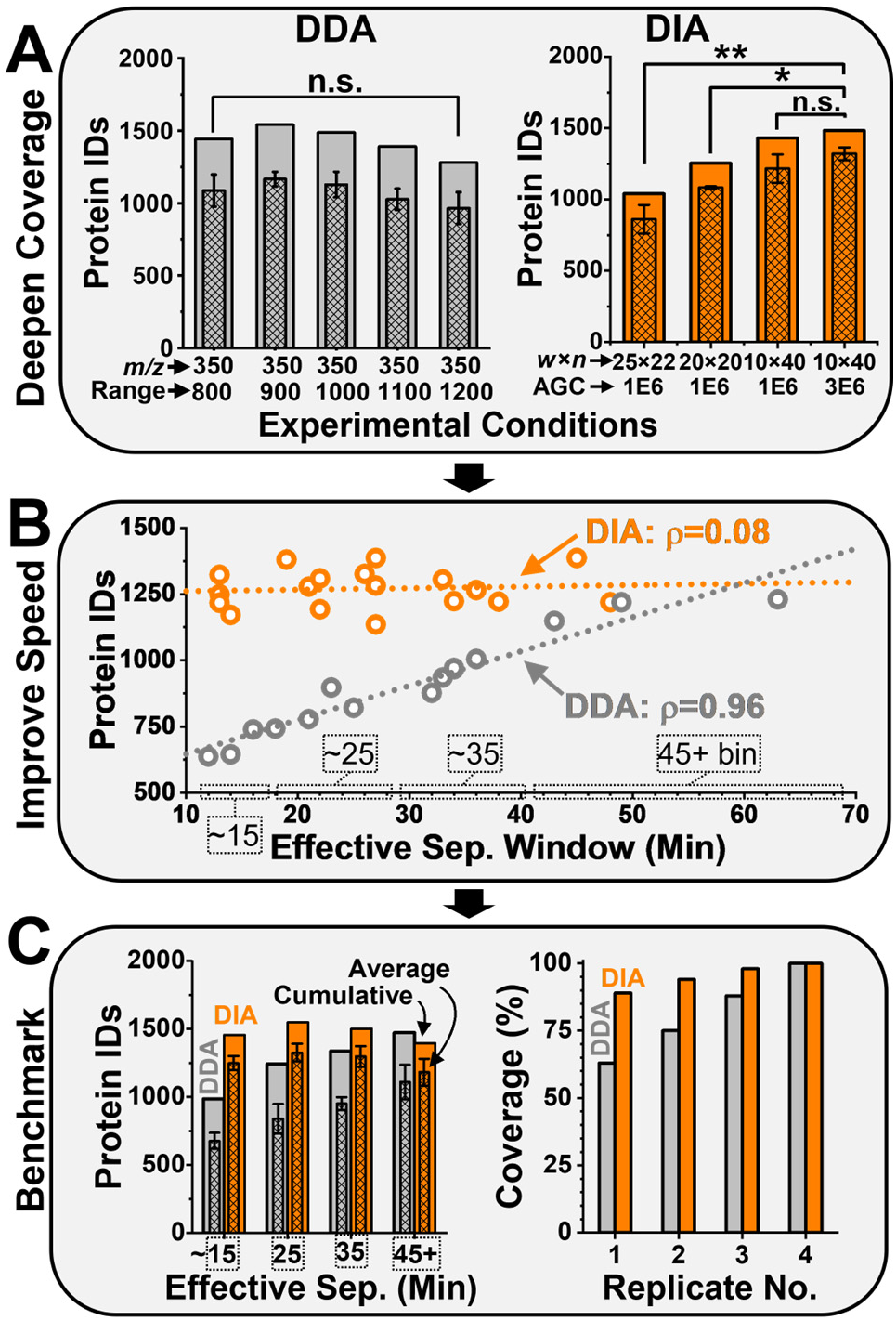
Development and benchmarking of high-throughput CE-MS proteomics. Each analysis measured ∼10 ng of the HeLa proteome digest. **(A)** The detectable coverage of the proteome was deepened by systematically exploring the experimental parameter space for data-dependent acquisition (DDA) and data-independent acquisition (DIA). Key: *n*, number of DIA windows; protein IDs, number of identified proteins; w, width of DIA window. **(B)** Shorter electrophoreses sacrificed protein identification in DDA, but not in DIA. **(C)** Compared to the DDA reference, <15-min CE-MS with DIA identified more proteins (**left panel**) and improved the reproducibility of the proteome coverage among technical replicates (**right panel**). Identification of 1,499 total proteins among 4 independent experiments maximized the experimentally detectable coverage (100%) of the proteome on our Q-Exactive Plus mass spectrometer.

To remedy this limitation, we revised the approach to operating the mass spectrometer. We posited that data-independent analysis (DIA) would aid MS–MS/MS bandwidth utilization for CE, because peptides migrating through the capillary as transient, yet resolved peaks would theoretically be fragmented within a wide m/z-window rolling over the entire mass-to-charge (m/z) range^[22, 26]^. As such, we reasoned DIA to be able to identify proteins from the lower concentration domain of the proteome compared to the standard DDA method, which prioritizes precursor peptide ions of higher ion signal abundance (concentration) for sequencing. In a series of pilot experiments (data not shown), we found the scanned m/z range, DIA m/z bin size × m/z window width, and ion accumulation time to notably impact the success of identifications. As shown in **Figure 2A**, the width and number of isolation windows and the number of ions accumulated (automatic gain control, AGC) mutually affected performance under the same C-trap ion accumulation time (55 ms). A narrower isolation width (10 m/z) reduced the complexity of chimeric tandem mass spectra and accumulation of more ions (3×10^6^ counts) enhanced detection sensitivity. The optimal conditions yielded 1,483 proteins from 10 ng of HeLa proteome digest within ca. 50 min effective separation duration on a previous-generation orbitrap mass spectrometer (Q-Exactive Plus, **SI Methods**).

### Our next objective was to speed up the measurements

We recognized electrical field strength in the BGE-filled CE capillary as a readily adjustable parameter to control the duration of electrophoresis. Facile and rapid conditioning of the CE capillary, e.g., by simple rinsing over <30 s after each measurement, provided another source of net gain in throughput. In comparison, the stationary phase in nanoLC typically requires relatively long equilibration times (e.g., 15 min to an hour), giving rise to the development of multi-column systems parallelizing separation and conditioning to remedy throughput.^[27]^ Indeed, the total analysis in our CE approach was finished in ∼105 min with a 65-min effective separation window at a low +10 kV of potential difference (100 V/cm field strength). This was shortened to ∼27 min with a 12 min effective window at +28 kV potential (280 V/cm field strength). Representative base-peak electropherograms illustrate these separations in **Figure S1**. By controlling the field strength, we were therefore able to analyze ∼10 ng of HeLa proteome digest over a series of separation times controlled over a broad range of duration.

Fast separation called for fast data acquisition. We implemented DIA with DDA serving as the benchmark on the same quadrupole-orbitrap mass spectrometer (Q-Exactive Plus, Thermo). The DIA library was developed by analyzing the HeLa proteome digest in 7 technical replicates using DDA, each time measuring ∼10 ng of the digest. **Figure 2B** presents the proteome coverage for effective separation windows ranging from ∼12 min to ∼65 min. The success of identifications progressively and rapidly decreased with shorter electrophoreses during DDA, but not DIA (library-based). For example, when employing the standard ∼50-min effective separation window, CE-MS identified 1,110 proteins on average (1,473 cumulative) using DDA, whereas DIA returned 1,183 proteins on average (1,396 cumulative) among 4 technical replicates. In stark contrast, a 15-min electrophoretic window produced only 677 proteins on average (986 cumulative) using DDA, compared to 1,249 proteins on average (1,457 cumulative) by DIA. Protein identifications are tabulated in **Tables S2–S5** for DDA and **Tables S6–S9** for DIA. **Figure 2C** bins the number of identified proteins as a function of the effective separation window (**left panel**) and explores the proteome overlap among the replicates for 15-min electrophoresis (**right panel**). Shorter separation times impaired proteome detection in DDA, whereas protein numbers held steady in DIA. Unlike in DDA where up to four replicate analyses still provided complementary proteome content, repeated measurements were unnecessary using DIA. At a short ∼15-min CE analysis time, library-based DIA doubled the coverage of the proteome compared to DDA. With ∼56% proteome shared by the approaches and DIA returning ca. 5-times more unique identifications, the library-based DIA outperformed DDA (**Fig. S2**). At 1,499 proteins identified, ∼15-min CE-MS (DIA, library-based) marked complete (100%) coverage of the proteome that was detectable using our Q-Exactive Plus mass spectrometer.

Library-free DIA promised additional speed, convenience, and scalability for limited proteomes by eliminating the need to experimentally prepare the spectral library. An artificial neural network^[28]^ executing a library-free search returned 1,403 cumulative proteins from the 15-min separations. Compared to 1,457 proteins resulting from the library-based alternative, ca. 71% of the detected proteomes were shared by the DIA approaches, with either returning a similar number of unique protein identifications (**Fig. S3**). Therefore, we considered the library-free DIA most beneficial for CE-MS proteomics. Because the size and quality of the spectral library affects protein identification and quantification,^[29]^ we also evaluated the quantitative performance of CE-MS (DIA, library-free) against DDA using calculated label-free quantification (LFQ) indices to approximate concentration.^[10]^ Indeed, 3 out of 4 technical replicates yielded quantitative information for ∼81% of the detected proteome (795 proteins) using DDA, only ∼43% of which (426 proteins) exhibited ≤20% of coefficient of variation (CV) based on LFQ (**Table S10**). In contrast, ∼89% of the measured proteome (1,343 proteins) was repeatedly quantifiable using DIA (library-free), ∼76% of which (1,073 proteins) had ≤20% CV (**Table S11**). Therefore, library-free DIA (termed DIA from here on) not only sped up protein analyses by eliminating the need for experimental library preparation *a priori* but also improved the depth and fidelity of quantification.

These performance differences indicated mechanistic differences between the data acquisition approaches. **Figure 3** evaluates the peptide groups that were identified within 15-min CE-MS using DDA and DIA. Based on our working hypothesis, broad m/z isolation in DIA was anticipated to fragment multiple precursor ions, thus democratizing the sampling of ion signal abundance over a broader dynamic range of protein concentration in relation to DDA. We tested these scenarios on 10 ng of the HeLa proteome digest. Indeed, as shown in **Figure 3A**, DIA consistently identified a higher number of peptide groups throughout the duration of separation, returning in fact many signals that were only detectable using DIA. These “DIA-exclusive” peptide signals populated the lower domain of the calculated label-free quantification (LFQ) indices estimating concentration, with this enhancement being statistically significant (**Fig. 3B**, *p* = 2.82×10^−4^, Mann-Whitney test).

**Figure 3.**
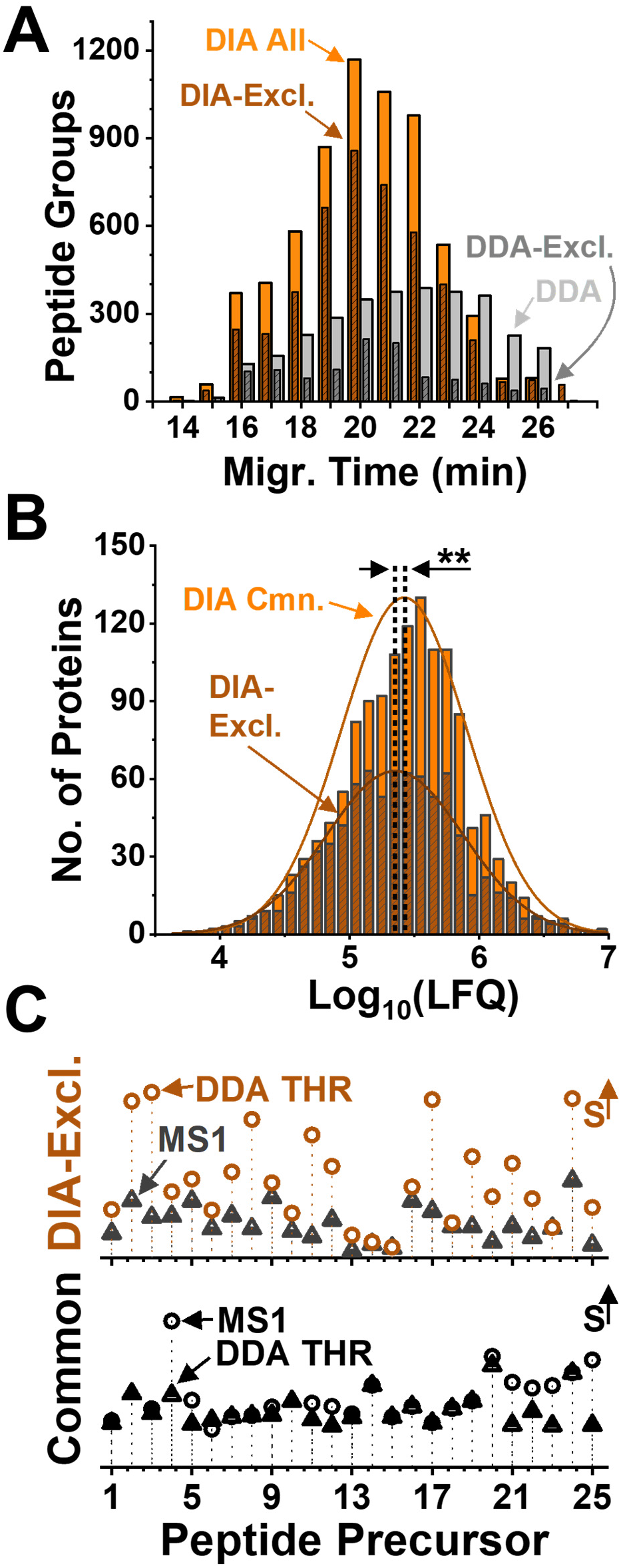
Mechanistic evaluation of DDA and DIA for CE-MS proteomics. As the standard, ∼10 ng of the HeLa proteome digest was measured in technical triplicates. **(A)** Comparison of the identification rate over electrophoresis time, revealing faster identifications by DIA vs. DDA **(B)** Proteins that were quantified exclusively by DIA were significantly lower in abundance than those shared by DDA (*p* = 2.82×10^−4^, Mann-Whitney). Key: **, p < 0.05. **(C)** Comparison of signal abundance (S) for the precursor peptide (MS^1^, data in circles) and their lowest threshold triggering (THR) MS/MS in DDA (data in triangles) for 25 randomly selected peptides that were **(top panel)** identified exclusively by DIA (DIA-Excl.) and **(bottom panel)** mutually with DDA.

We explored the underlying mechanism of the observed sensitivity gains. **Figure 3C** tests our working hypothesis predicting sensitivity improvement for lower-concentration signals due to lacking signal thresholding in DIA. We compared in the DDA measurements the precursor (MS^1^) signal abundance of 25 randomly selected, DIA-exclusive peptides (accurate m/z values ± 5 ppm error) against the weakest peptide ion signal in the top-10 train of tandem (MS^2^) transitions (**top panel**). Indeed, these the DIA-exclusive peptides did not reach sufficient threshold for consideration for MS/MS in DDA. For a group of 25 different peptide signals that were mutually identified by DDA and DIA despite producing low ion signal abundance, the criteria of high-pass filtering and top-10 prioritization were met for tandem MS by DDA. Surprisingly, <4% of the quantified proteome were measured exclusively using DDA. The proteotypic peptide signals corresponding to these identifications populated the entire concentration range of the proteome (DDA-Excl. vs. DDA-All: *p* = 0.71, Mann-Whitney test). Approximately 60% of these peptides had m/z values outside the optimized m/z range of DIA; these peptides were not dissociated by default. For 25 randomly selected peptide signals that were exclusively detected by DDA within the DIA m/z range, 21 were detected in the DIA MS^1^ spectra. Closer inspection of the primary MS-MS/MS data revealed that elongated DIA cycle times missed the transient signal for 4 of these peptides during wide m/z-window fragmentation. Challenges associated with the processing of complex chimeric mass spectra may account for the other peptide ions. Combined, these mechanistic insights confirmed superior sensitivity and quantitative reproducibility from DIA compared to the DDA standard using <15-min CE-MS.

### These performance improvements raised the potential for single-cell proteomics in <15-min

As severely limited proteomes, such as mammalian (HeLa) cells, give inherently less ion signal abundance, we proposed to further detection sensitivity by elongating the ion accumulation time on both the survey (MS^1^) and tandem MS (MS^2^) scans. To test scalability to single mammalian cells, we electrophoresed in <15 min an ∼200 pg portion of the digest, which estimates to the proteome content of a single HeLa cell. DIA executing the scheme of 10 windows × 40 m/z width/window with accumulation times of 240 ms for MS^1^ and 50 ms for MS^2^ returned 474, 502, 519, 549, and 528 proteins among the technical replicates (579 cumulative), which amounted to 974 proteins among the files using the match-between run (MBR) algorithm (**Table S12**). In contrast, DIA executing the 20 × 20 scheme with 240 ms accumulation time for MS^1^ and 110 ms for MS^2^ returned 653, 547, 569, 543, 485 protein identifications among the replicates (684 cumulative), totaling to 1,161 proteins using MBR (**Table S13**). Therefore, longer ion accumulation times deepened CE-MS (DIA) coverage. These results mark a substantial sensitivity improvement over the reference DDA approach, which found 401 proteins (data not shown). In comparison, recent analyses of the HeLa proteome on similar-generation orbitrap instruments as used here detected ∼700 proteins over ∼120 min of gradient using LC^[30]^ and ∼1,100 proteins from 880 pg in ∼50 min of effective separation window by a commercial CE instrument.^[14]^ Therefore, longer accumulation times notably improved the detectable proteome using ∼15-min CE-MS (DIA).

With spatiotemporally scalable capillary microsampling, CE-MS opened the door to deep subcellular proteomics. **Figure 4** presents this portion of the study. As the biological model, we selected the midline dorsal–animal (called D11) embryonic cell in the 16-cell live *X. laevis* embryo. With a considerable, ∼250-μm-dimeter size (∼90 nL volume) and a short, ∼15-min (room temperature) life span before division, we microfabricated borosilicate capillaries following our recent studies^[12]^ to aspirate ∼10 nL contents of the cell from randomized locations (yielding ∼167 ng of yolk-free proteome). Each of the collected proteomes was trypsin-digested and reconstituted to 5 μL in low-bind Eppendorf vials (see **SI** Materials and Methods), and ∼5 ng of each proteome sample was measured in 15 min using CE-MS executing DIA and DDA (reference). Each experiment was repeated N = 5 times using different embryos (biological replicates), with each proteome sample analyzed in 3 technical replicates. The total number of proteins that were identified from the subcellular aspirates were 825 by DDA (**Table S14**) and 1,242 by DIA (**Table S15**). While the control DDA gave a comparable number of proteins to our previous work, demonstrating technical robustness^[12]^, the DIA approach identified ∼1.5-times more proteins. In comparison to recent nanoLC^[5c]^ and CE^[31]^ analyses employing a comparable generation of mass spectrometer to that used here, CE-MS (DIA) accelerated measurement throughout by ∼4.5–8-fold and deepened detection sensitivity by ∼8–10-times.

**Figure 4.**
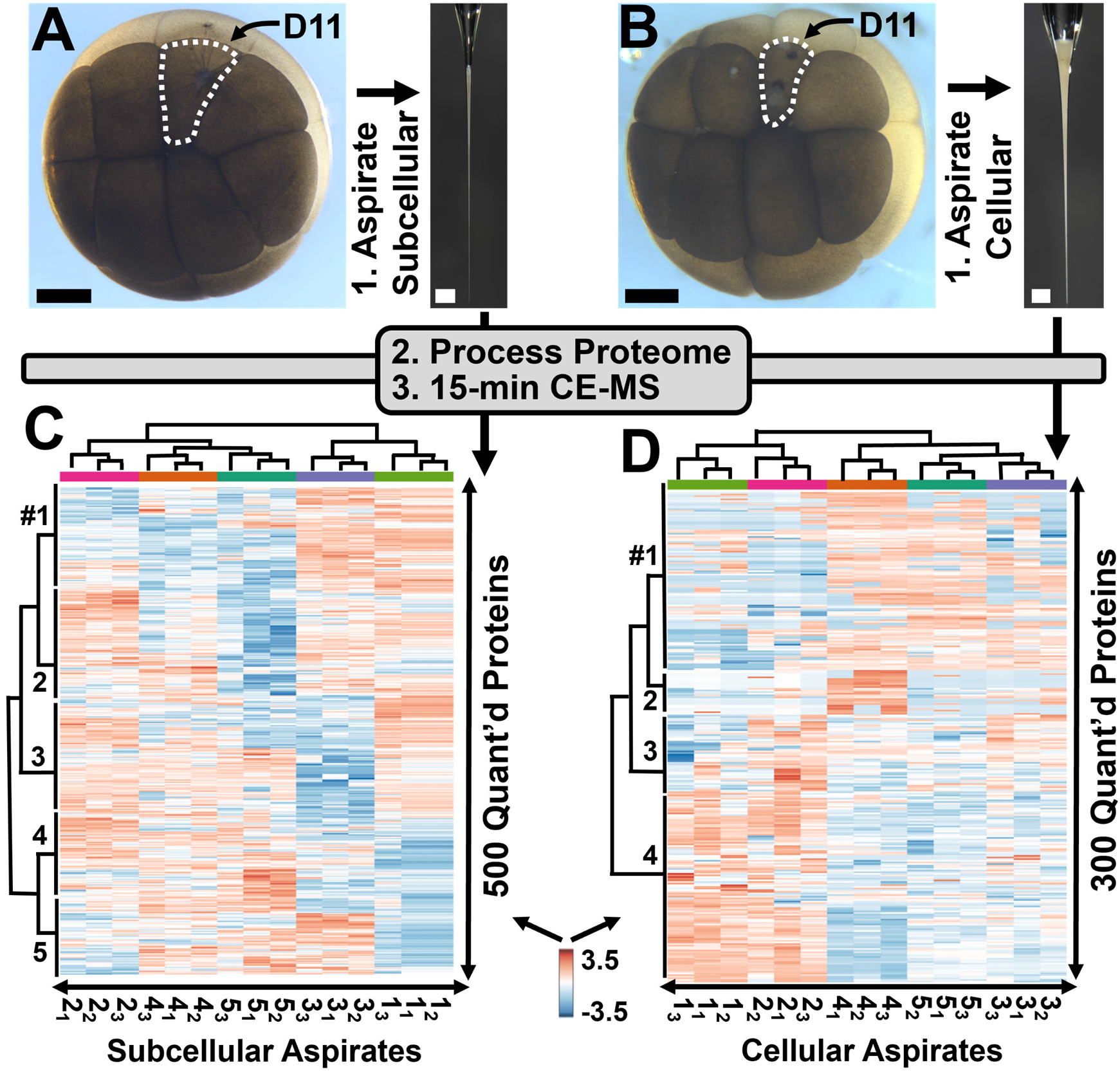
15-min proteome analysis on the midline dorsal-animal cell in live 16-cell *X. laevis* embryos. Using a fabricated microcapillary, **(A)** a 10 nL portion of the cell and **(B)** the intracellular ∼90 nL content of the identified cell was aspirated in N = 5 biological replicates (each from a different embryo). The collected proteomes were quantified by CE-MS (DIA) with a <15 min effective separation window. Hierarchical cluster–heat map analysis of the **(C)** 500 and **(D)** 300 most significantly differently expressed proteins revealed subcellular/cellular proteomic differences. Labels #1–5 mark proteins with similar abundance profiles among the samples. Key to each replicate: Biological_Technical_. Scale bars, 250 μm (black) and 300 μm (white).

Gene ontology annotation provided insights on the canonical function of the newfound proteins. **Table S16** compares the proteins that were identified between the subcellular aspirates from the D11 cells using DDA and DIA. Annotation of the DIA-exclusive data in PantherDB version 18.0^[32]^ assigned 143 proteins to intracellular organelles, such as the mitochondrion, nucleus, endoplasmic reticulum, and Golgi apparatus. Detection of the organelle-affiliated proteins validated the success of subcellular sampling. The detected proteins are known to serve important canonical functions. For example, 93 and 91 proteins fulfill binding and catalytic activity, respectively. Proteins identified exclusively by DIA are involved in a broad range of biological processes, including cellular metabolic processes (76 proteins), biological regulation (44 proteins), and localization (27 proteins). Therefore, capillary microsampling with CE-MS (DIA) obtained previously inaccessible proteomic information on the subcellular organization of the cell.

With LFQ, the approach offered a key to profiling proteome differences between the cells and their contents. In theory, we anticipated randomized microprobe sampling to be sensitive to subcellular proteome niches, differences in cell cycle, and/or natural proteomic variability between the embryos. To help decipher these scenarios, we also analyzed the entire intracellular proteome from N = 5 different D11 cells using a microcapillary (**Fig. 4B**), excluding the outer membrane of the cells. We expected these quasi-cellular aspirates to reflect information on cell cycle and embryo variability. The calculated LFQ abundance indexes were log_10_-transformed and mean normalized, thus allowing us to compare equal amounts of proteomes. **Figure 4** presents unsupervised multivariate analysis for the most differentially expressed proteins on the subcellular (**panel A**) and quasi-cellular analyses (**panel B**). Based on their concentration variations among the aspirates, the hierarchical cluster analysis (HCA)–heat map grouped the proteins into 5 subclusters in the subcellular data (labels #1*–*5 in **Fig. 4C**) and 4 subclusters in the cellular (labels #1–4 in **Fig. 4D**). STRING version 12^[33]^ interpretation of subcellular localization and molecular roles appreciated differences between the subcellular data, but not in the cellular analyses. In the subcellular aspirates, 67% of the proteins in cluster 1 were enriched mitochondria and related processes, 17% of the proteins in cluster 2 were annotated with the ribosome and ribosomal activity, and 40% of the detected proteins were affiliated with the nucleus and nuclear processes in cluster 3 (**Table S17**). In contrast, a similar HCA analysis on the quasi-cellular proteomes revealed comparable organelle and molecular composition, suggesting that the subclusters are due to differences in cell cycle or natural biological variability among the cells (**Table S17**). Combined, these results demonstrate the sensitivity and fidelity of CE-MS (DIA) to characterize proteome differences between and within cells.

## CONCLUSIONS

This study enabled the 15-min deep proteome analysis of single cells and their contents. We demonstrated that CE is capable of fast and reproducible separation for complex proteome digests, with rapid (<30 s) rinsing of the capillary further elevating measurement throughput. Compared to classical DDA, we found DIA to alleviate limitations in the duty cycle of tandem MS and democratized molecular coverage across the concentration range of the endogenous proteomes. From single-HeLa-cell-equivalent proteomes as the benchmark, the approach returned 1,161 proteins on an early-generation Orbitrap mass spectrometer (Q-Exactive Plus), nearly tripling proteome coverage compared to the DDA standard. With spatiotemporally scalable microprobe sampling, this sensitivity was sufficient to measure 1,242 proteins and distinguish subcellular environments in single identified cells (D11) in live *X. laevis* embryos, including several canonical protein components of organelles. These identifications and analytical speeds mark an 8–10-fold improvement in sensitivity and 4.5–8-fold enhancement in throughput compared to LC-MS studies that used a comparable generation and configuration of mass spectrometer. Although analyte losses on vial surfaces inherently hindered molecular coverage in the current work, we expect recent technologies enabling the processing of limited proteomes (e.g., nanoPOTs^[34]^ and mPOP^[35]^) to be readily adoptable to deepen the detectable proteome. Of course, the use of late-generation mass spectrometers with substantially higher sensitivity and faster data acquisition (e.g., Thermo Orbitrap Astral and Bruker timsTOF Ultra) are bound to further CE-ESI sensitivity to ever-deeper coverage of the cellular proteome in high throughput. CE-MS with DIA expands the bioanalytical toolbox of cell and molecular biology to the cell and its subcellular environments in scalability over space, time, and molecular coverage.

## Acknowledgment

We thank Kellen DeLaney (University of Maryland, College Park) for discussions on general concepts of DIA. Parts of this research were supported by the National Institute of General Medical Sciences (award no. R35GM124755 to P.N.), the Arnold and Mabel Beckman Foundation (Beckman Young Investigator Award to P.N.), and the University of Maryland Brain and Behavior Institute (seed award to P.N.).

## Data Repository

All MS–MS/MS primary files and the MS/MS spectral libraries, and the HeLa and *Xenopus* proteomes were deposited to the ProteomeExchange Consortium via the PRIDE partner repository with the data set identifier PXD046467.

