## Supplementary Document for "The 15-min (Sub)Cellular Proteome"

### MATERIALS AND METHODS

**Chemicals and Materials.** All the solvents and chemicals including acetic acid (AcOH), acetonitrile (ACN), formic acid (FA), and methanol (MeOH) were purchased from Fisher Scientific at LC-MS grade. Ammonium bicarbonate (AmBic) was from Avantor (Center Valley, PA). The HeLa proteome digest standard was supplied by Thermo Fisher (part no. 88329, Pierce, Rockford, IL). Microcapillaries for subcellular sampling and CE-ESI were fabricated from borosilicate glass capillaries (0.75/1.00 mm inner/outer diameter, part no. B100-75-10 and 0.50/1.00 mm inner/outer diameter, part no. B100-50-10, Sutter Instrument, Novato, CA, respectively) using a micropipette puller (P-1000, Sutter Instrument, Novato, CA). For CE, fused silica capillaries were obtained from Polymicro Technologies (40/105 μm inner/outer diameter, part no. 1068150596, Phoenix, AZ) and used as received. Proteins were digested with MS-grade TPCK-modified trypsin (part no. 90057, Pierce, Rockford, IL). All protein and peptide samples were processed in 0.2-mL LoBind Eppendorf vials to reduce molecular losses due to nonspecific adsorption on container surfaces (part no. 951010006, Eppendorf, USA).

**Solutions.** The contents of single cells were aspirated into 50 mM AmBic prepared with HPLC water. The CE *background electrolyte* (BGE) consisted of 25% (v/v) ACN in water with 1 M FA. The CE-ESI *sheath solution* contained 10% (v/v) MeOH in water with 0.5% (v/v) AcOH. The *sample solvent* contained 75% (v/v) ACN in water with 0.05% (v/v) FA.

**Animal Care and Embryology.** Adult, sexually mature *Xenopus laevis* frogs were received from Nasco (Fort Atkinson, WI) or Xenopus1 (Dexter, MI). All protocols regarding handling of *X. laevis* were approved by the Institutional Animal Care and Use Committee of the University of Maryland, College Park (Approval no. #R-FEB-21-07). Embryos were obtained from natural mating of multiple pairs of parents to account for natural variability. Two-cell embryos displaying stereotypical pigmentation were selected and cultured to the 16-cell stage following standard protocols.[1] The left dorsal-animal midline (called D11) cell was identified based on location, morphology, pigmentation, and in reference to reproducible fate maps.[1]

**Subcellular Proteome Sampling.** We performed capillary microsampling to collect a portion of the single-cell proteome. As described elsewhere,[2] a borosilicate capillary was pulled to ~20-μm-diameter tip and mounted on a three-axis translation stage to pierce an identified D11 cell. A pulse of negative pressure was applied to the back end of the capillary to aspirate ~10 nL portion of the cell content. Using a positive pressure pulse, the collected cell contents were expelled into a 0.2-mL LoBind microtube containing 5 μL of 50 mM AmBic solution. The conventional bottom-up proteomics steps of protein reduction and alkylation were eliminated to minimize sample loss and improve protein identification.[2-3] Instead, the subcellular proteome was denatured by rapid heating to 60 ºC for 20 min, before digestion to peptides with the addition of 1 μL of 0.25 μg/μL trypsin and incubation at 37 ºC for 5 h. The resulting peptide mixture was vacuum-dried at 60 ºC and stored at –80 ºC until analysis.

**CE-nanoESI-HRMS.** The samples were measured on a CE-nanoESI platform that we custom-built following our earlier designs.[2, 3b] In this study, ~10 ng of HeLa proteome digest were hydrodynamically injected into a ~100-cm-long CE separation capillary that was filled with the BGE. Electrophoresis was performed by applying a constant potential to the inlet end of the CE separation capillary as follows: +28 kV (for ~30 min separation), +25 kV (for ~40 min separation), +20 kV (for ~60 min separation), +15 kV (for ~75 min separation), or +12 kV (for

+90 min separation) against Earth ground. The outlet end of the CE separation capillary was fed into an electrokinetically pumped sheath-flow CE-nanoflow ESI interface that we custom-built following our earlier design.[2] The CE-nanoESI interface was mounted on a three-axis translation stage to position the tip of the electrospray emitter ~1 mm from the orifice of a mass spectrometer. At +900–1,200 V applied to the inlet end of the electrokinetic pump, the electrospray was maintained in the cone-jet regime, thus maximizing the efficacy of ion production.[4] The spraying regime was identified based on the electrohydrodynamic behavior of the Taylor cone, monitored by long-working distance objective microscope (Mitutoyo Plan Apo, Edmund Optics, Barrington, NJ) with a CCD camera (EO-2018C, Edmund Optics).

The generated ions were analyzed using a quadrupole orbitrap mass spectrometer (Q-Exactive Plus, Thermo Scientific) executing data-dependent acquisition (DDA) or data-independent acquisition (DIA) for tandem MS. In both data acquisition modalities, the ion signals (m/z values) were dissociated in nitrogen collision gas at 28% normalized collision energy (NCE) in the higher-energy collisional dissociation (HCD) cell. The specifics of each data acquisition approach are detailed in the following.

***DDA.*** Survey $MS^1$ scans were acquired using the following settings: Orbitrap mass resolution, 70,000 full width at half max (FWHM at *m/z* 200); scan ranges, *m/z* windows (350–800; 350–900; 350–1,000, 350–1,100; and 350–1,200); C-trap maximum injection time, 50 ms; AGC target, $1 \times 10^6$ counts. The 10 most abundant ("top") precursor ions were selected for fragmentation using the following settings: m/z isolation window, 1.5 m/z; mass resolution, 17,500 FHWM (*m/z* 200); C-trap maximum injection time, 100 ms; and AGC target, $5 \times 10^4$ counts.

***DIA.*** The DIA mass isolation windows were generated in Skyline[5]. The "optimizing window placement" method was executed to avoid placing edges of the isolation windows in regions where peptides were likely to occur between consecutive mass windows. In this study, 4 different DIA conditions were tested, which are summarized in the following combinations of m/z scan range (low–high m/z), window schemes (window width × number of windows), and AGC target (counts), respectively: 350–900, 22 × 25, and $1 \times 10^6$; 490–910, 20 × 20, and $1 \times 10^6$; 490–910, 10 × 40, and $1 \times 10^6$; 490–910, 10 × 40, and $3 \times 10^6$. The following $MS^1$ parameters were used: Orbitrap mass resolution, 35,000 FWHM (*m/z* 200); C-trap maximum injection time, 55 ms. $MS^2$ parameters were as follows: mass resolution, 17,500 FHWM (*m/z* 200); C-trap maximum injection time, auto.

**Benchmarking to a Single HeLa-Cell-Equivalent Proteome Amount.** The performance of CE-ESI-MS was assessed for DIA by analyzing 200 pg of standard HeLa proteome digest, which approximates to the total proteome content of a single HeLa cell.[6] For these trace amounts of proteomes, the C-trap injection time was increased to a maximum of 240 ms. The $MS^2$ scans employed the following combinations of mass window (window width × number of windows), Orbitrap mass resolution, maximum C-trap injection time, and maximum AGC target: 20 × 20, 35,000 FWHM, 110 ms, and $3 \times 10^6$ counts; and 10 × 40, 17,500 FWHM, 50 ms, and $3 \times 10^6$ counts.

**Data Analysis**. ***DDA***. The MS–MS/MS data were analyzed in Proteome Discoverer version 2.2 (Thermo) on the SEQUEST search engine with the minora algorithm for LFQ. For the HeLa proteome digest, identifications were compared against the UniProt Human Proteome Database (UP000005640, downloaded on June 2021, containing 20,380 entries). The search parameters were the following: static modification, cysteine carbamidomethylation; dynamic modifications,

methionine oxidation; minimum peptide length, 5 amino acids; maximal missed cleavage sites, 2; precursor and fragment ion mass tolerance, 10 ppm and 0.02 Da, respectively; search for common contaminants enabled. For the *X. laevis* subcellular proteome, the MS–MS/MS data were mapped against the *Xenopus laevis* UniProt Proteome Database (UP000694892, downloaded on April 2022, containing 42,596 entries). The following search settings were employed: static and dynamic modifications, excluded (none used); precursor and fragment ion mass tolerance, 10 ppm and 0.02 Da, respectively. For the technical replicates, the total peptide amount normalization was performed. Protein identifications were made based on the detection of at least 1 proteotypic peptide. Peptides and protein identifications from HeLa or *X. laevis* were filtered to <1% false discovery rate (FDR) against a reversed-sequence decoy database.
***DIA***. The DIA MS–MS/MS data were processed in Spectronaut version 15 (Biognosys Inc., Cambridge, MA). The DIA MS/MS spectra from 10 ng of HeLa proteome digest were searched against a spectral library that was custom-built by measuring 10 ng of HeLa proteome digest using DDA in 7 technical replicates. The DIA MS/MS data from the *X. laevis* subcellular proteome and the single-cell-equivalent of HeLa proteome digest (200 pg) were searched library-free between 3–5 technical replicates of each sample using DIA-NN[7] with the following settings: match between runs, enabled; minimal peptide length, 5; maximal peptide length, 35; maximum missed cleavage, 2; maximum number of oxidation as variable modification, 1; and precursor charge, 2–4. All the other parameters were set to default. Protein identifications were made based on the detection of at least 1 proteotypic peptide. Peptides and protein identifications from HeLa or *X. laevis* were filtered to <1% false discovery rate (FDR) against a reversed-sequence decoy database.
**Scientific Rigor.** The HeLa proteome digest was measured in 3–5 technical replicates (same sample analyzed multiple times). *X. laevis* subcellular proteome was measured in 5 biological replicates (each from a different embryo), with each biological replicate analyzed in 3 technical replicates.

**Safety.** All chemicals and biological samples were handled following standard safety protocols. Care was taken during the handling of capillaries and ESI emitters to mitigate potential puncture hazards. All electrically conductive components of the CE-HRMS platform were grounded or shielded from exposure in an enclosure with a safety interlock to avoid potential electroshock hazard.

### TABLES

**Table 1.** Comparison of protein identifications at different durations of CE separation using DDA and DIA methods. The resulting average and cumulative (∑) identifications are tabulated for each individual measurement on the same proteome sample.

| DDA | | | |
|---|---|---|---|
| Eff. Sep. (Min) | MT (Min) | IDs | Results Aver./∑ |
| 12 | 27 | 637 | |
| 14 | 26 | 645 | 691 ± 58 |
| 16 | 29 | 739 | 974 |
| 18 | 31 | 742 | |
| 21 | 39 | 778 | 832 ± 60 |
| 23 | 41 | 897 | 1,069 |
| 25 | 40 | 821 | |
| 32 | 57 | 877 | |
| 33 | 56 | 935 | 937 ± 43 |
| 34 | 56 | 966 | 1,266 |
| 34 | 55 | 971 | |
| 36 | 60 | 1,005 | |
| 43 | 86 | 1,148 | 1,199 ± 45 |
| 49 | 86 | 1,219 | 1,549 |
| 63 | 120 | 1,230 | |

| DIA | | | |
|---|---|---|---|
| Eff. Sep. (Min) | MT (Min) | IDs | Results |
| 13 | 28 | 1,249 | |
| 13 | 27 | 1,323 | 1,254 ± 63 |
| 13 | 29 | 1,271 | 1,495 |
| 14 | 28 | 1,171 | |
| 19 | 39 | 1,380 | |
| 21 | 37 | 1,278 | 1,290 ± 77 |
| 22 | 43 | 1,193 | 1,656 |
| 22 | 43 | 1,309 | |
| 26 | 51 | 1,326 | |
| 27 | 53 | 1,282 | |
| 27 | 52 | 1,135 | 1,276 ± 87 |
| 27 | 52 | 1,386 | 1,631 |
| 33 | 61 | 1,305 | |
| 34 | 63 | 1,223 | |
| 36 | 81 | 1,265 | |
| 38 | 77 | 1,221 | 1,273 ± 78 |
| 45 | 92 | 1,386 | 1,680 |
| 48 | 75 | 1,219 | |

### FIGURES

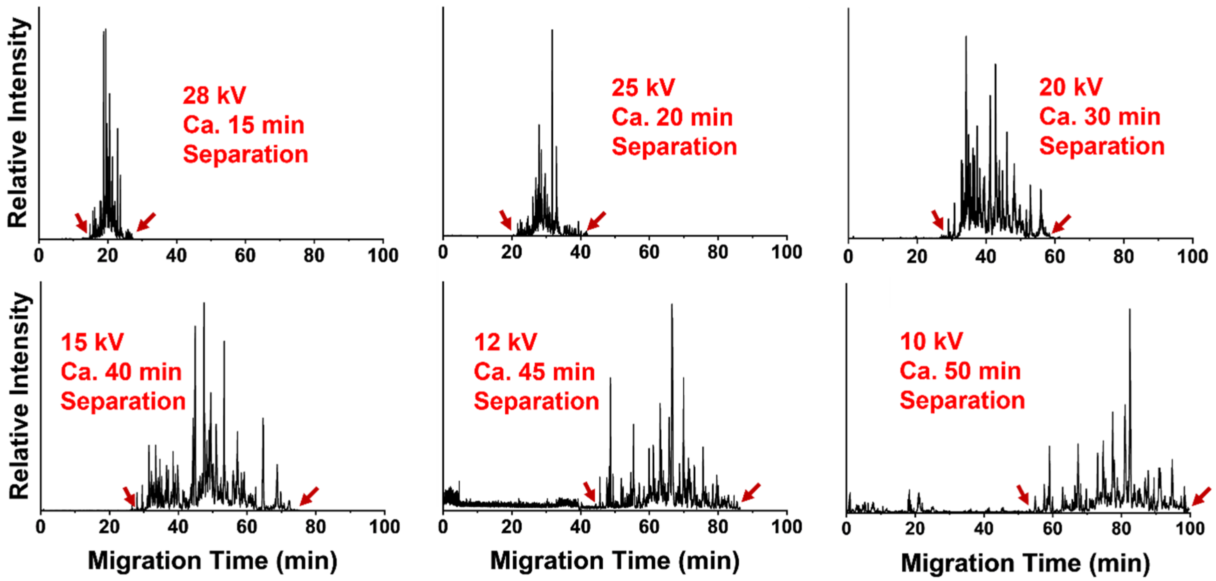


**Figure S1.** Base-peak electropherograms showing adjustable separation duration. A total of ~10 ng of the HeLa proteome digest was analyzed in each experiment. The potential difference was adjusted between the inlet and outlet ends of the background-electrolyte filled CE capillary to control separation time. The arrows mark the begin and end of each effective separation window.

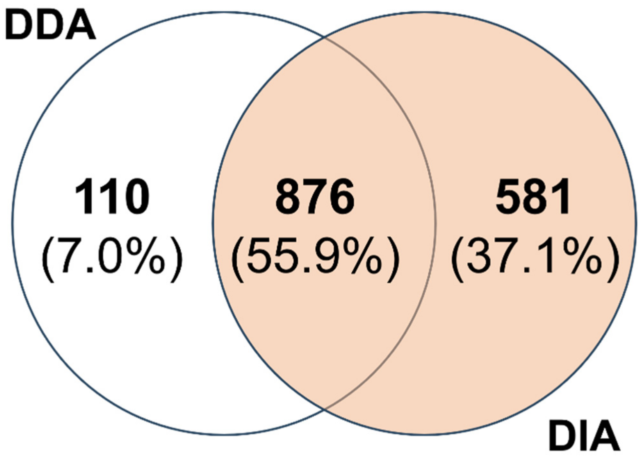


**Figure S2.** Comparision of protein identification from N = 4 technical replicate measurement of 10 ng of the HeLa proteome digest in each using 15-min CE-MS with DDA and DIA (library-based). The library-based DIA approach outperformed the DDA.

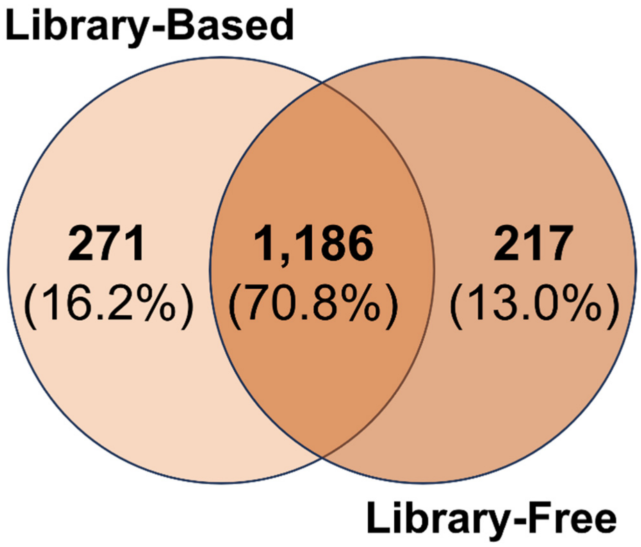


**Figure S3.** Comparision of the number of proteins that were identified between library-based and library-free CE-MS (DIA). We considered the performance of the approaches similar. For limited amounts of proteomes, the library-free approached eliminated the need for experimental measurement of the library using DDA.
